# Polarization of microbial communities between competitive and cooperative metabolism

**DOI:** 10.1101/2020.01.28.922583

**Authors:** Daniel Machado, Oleksandr M. Maistrenko, Sergej Andrejev, Yongkyu Kim, Peer Bork, Kaustubh R. Patil, Kiran R. Patil

**Affiliations:** European Molecular Biology Laboratory (EMBL), Meyerhofstraße 1, 69117 Heidelberg, Germany; Institute of Neuroscience and Medicine, Brain and Behaviour (INM-7), Research Center Jülich, Jülich, Germany; Institute of Systems Neuroscience, Medical Faculty, Heinrich Heine University Düsseldorf, Düsseldorf, Germany; MRC Toxicology Unit, University of Cambridge, Cambridge, UK

## Abstract

Resource competition and metabolic cross-feeding are among the main drivers of microbial community assembly. Yet, the degree to which these two conflicting forces are reflected in the composition of natural communities has not been systematically investigated. Here, we use genome-scale metabolic modeling to assess resource competition and metabolic cooperation potential in large co-occurring groups, with up to 40 member species, across thousands of habitats. Our analysis revealed two distinct community types, clustering at opposite ends in a trade-off landscape between competition and cooperation. On one end lie highly cooperative communities, characterized by smaller genomes and multiple auxotrophies, reminiscent of the black queen hypothesis. At the other end lie highly competitive communities, conforming to the red queen hypothesis, featuring larger genomes and overlapping nutritional requirements. While the latter are mainly present in soils, the former are found both in free-living and host-associated habitats. Community-scale flux simulations showed that, while the competitive communities can better resist species invasion but not nutrient shift, the cooperative communities are susceptible to species invasion but resilient to nutrient change. In accord, we show, through analyzing an additional independent dataset, the colonization of the human gut by probiotic species is positively associated with the presence of cooperative species in the recipient microbiome. Together, our analysis highlights the bifurcation between competition and cooperation in the assembly of natural communities and its implications for community modulation.

## Introduction

Microbial communities are fundamental constituents of ecosystems across scales^1–6^. They play a crucial role in, for example, geochemical cycles^7^ and in our health as our microbial symbionts^7^. The biological properties and functions of these communities are determined by their compositional make-up. For example, multiple diseases have been linked to compositional changes in the gut microbiome^5,8^. An emerging challenge in these and other microbial ecosystems is modulation or redesign of communities towards repairing a perturbed state or reaching a new community-level function^9^. However, it is currently difficult to predict which microbes would form a stable community or how a given community would respond to different biotic or abiotic perturbations.

Nutrient availability is one of the most fundamental factors directing the establishment of a community and the metabolic interactions therein^10–12^. Previous studies have assessed the potential for such interactions using genome-scale metabolic models, which recapitulate the metabolic and biosynthetic capabilities of each community member. While these studies attest to the potential of genome-scale modeling to assess the competition and cooperation in microbial communities, they remain limited due to the low species numbers and habitat diversity analyzed. Furthermore, the restricted size of co-occurring groups (only species pairs) does not capture higher-order interactions (i.e. the influence of a third, or more, species in the interaction between two species), which are known to play a role in ecosystem function^13–15^.

Here, we assessed, through simulating thousands of community-scale metabolic models, the prevalence and the nature of metabolic competition and cross-feeding interactions in microbial communities across thousands of samples from diverse environments represented in the Earth Microbiome Project (EMP)^16^ and validated some of the derived hypothesis using additional independent datasets. The broad habitat coverage and consideration of large co-occurring groups allowed us to gauge the relative role of metabolic competition and cooperation in community assembly, and the evolutionary signatures of this trade-off in microbial genomes.

## Results

### Co-occurring communities

We first built metabolic models for individual species by mapping 16S rRNA sequences present in the EMP dataset, previously classified into operational taxonomic units (OTUs), to their closest reference genomes in NCBI RefSeq, a database of fully sequenced genomes^17^, using a 97% similarity cutoff (see Methods). We then used these genomes to build genome-scale metabolic models with CarveMe^18^. This resulted in a collection of unique models for 2986 species. Next, to uncover ecologically-relevant patterns of interactions among these species, we systematically searched for groups of significantly co-occurring species (i.e. groups of species that occur together across samples more often than expected by chance; see illustration in Figure 1a, and Methods for details). Although multiple methods have been proposed to compute co-occurring species in microbial samples, most are limited to species pairs^19–22^. On the other hand, experiments with synthetic communities have underlined the importance of higher-order interactions in community structure and dynamics. The emergent features of complex communities thereby cannot be inferred from pairwise interactions alone^15^. Supporting this, our previous work showed that co-occurring communities could be distinguished from random species assemblies much more markedly in triplets and quadruplets^23^.

**Figure 1.**
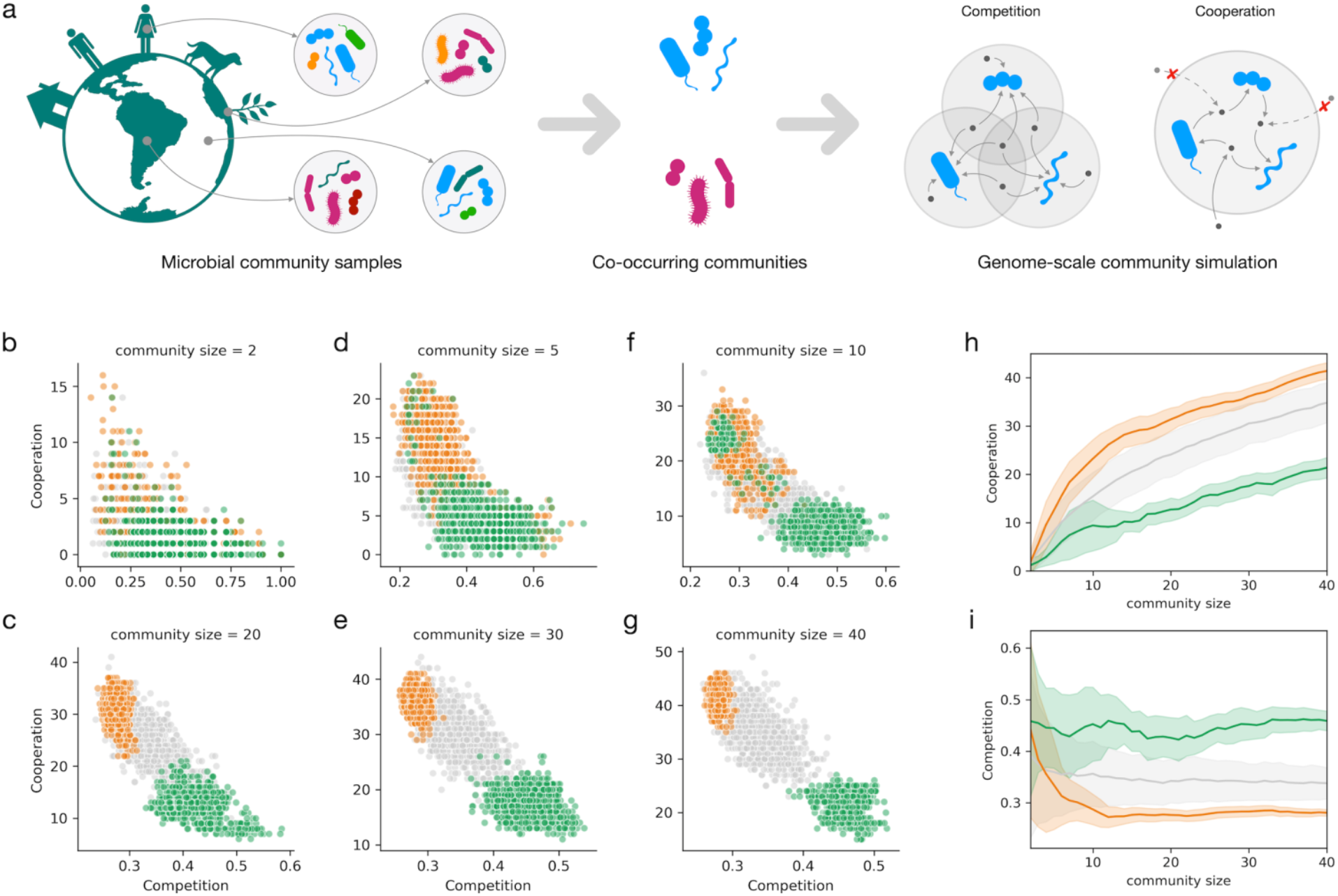
a) Schematic of the two main steps in our analysis: identification of frequently co-occurring communities across the Earth Microbiome project samples, followed by the calculation of metabolic resource overlap (MRO) and metabolic interaction potential (MIP) scores; b-g) The trade-off between the competition (MRO) and cooperation (MIP) scores for different community sizes. Green and orange dots show co-occurring communities, while the grey dots represent random assemblies, with a total of 1000 simulated communities per each community size and type; h) Cooperation potential as a function of community size; i) Competition potential as a function of community size (thick lines represent mean values and confidence intervals represent standard deviation).

In this work, we tackle higher-order interactions at an unprecedented scale by introducing a new heuristic approach (see Methods). In brief, the method begins by computing significantly cooccurring species pairs, and iteratively creates larger assemblies using a sampling approach, based on roulette wheel selection, to avoid the combinatorial explosion. This allowed us to uncover thousands of co-occurring communities with up to 40 member species. To visualize the distribution of species in these communities, we applied dimensionality reduction using principal component analysis, and observed the formation of two spatially segregated clusters of cooccurring communities (Supp. Fig. 2).

### Competition - cooperation tradeoff

We next assessed metabolic interactions in the identified co-occurring communities using SMETANA, a flux balance analysis-based simulation tool^24^. Unlike other community simulation methods^24–26^, SMETANA does not assume any optimality at community or species level, its only assumption is that each species can survive using the available resources. Using SMETANA, we computed the metabolic resource overlap (MRO) and metabolic interaction potential (MIP) for each community (as illustrated in Fig. 1a). The MRO quantifies the similarity of the nutritional requirements between all species in a community, reflecting the intra-community risk for resource competition. The MIP provides an estimate for the number of metabolites that can be exchanged among the community members to decrease their dependency on the nutrient supply from the (abiotic) environment. As control groups, to contrast against the co-occurring communities, random assemblies of the same size as the co-occurring communities were used (1000 communities for each community size).

The first notable pattern from the simulation results was a trade-off between competition and cooperation (Fig. 1b). Species within communities with higher cooperation potential thereby have less resource overlap and vice-versa. This pattern, although intuitive, had not been observed before, most likely due to the limited scale of the previous studies. To check if the observed tradeoff pattern results from any biases in the EMP data (such as the habitats covered, experimental protocols, or data processing pipelines), we computed co-occurring communities using an independent collection of 16S amplicon data compiled from multiple sources by Chaffron and coworkers^21^. Again, we observed a clear trade-off between competition and cooperation (Supp. Fig. 3).

When compared with the random assemblies, the co-occurring communities not only showed a striking distinction in terms of both competition risk and cooperation potential but also a clear polarization at the opposite ends of the competition-cooperation spectrum (Fig. 1b). Furthermore, in accordance with the ecological importance of higher-order interactions, the distinction of the co-occurring groups is more prominent for larger community sizes. The two polarized clusters of the co-occurring communities, the highly competitive and highly cooperative, coincide with the two main clusters previously observed based on species composition (Supp. Fig. 2). This stark contrast suggests opposite metabolic strategies undertaken by the species present in the two groups.

### Species characteristics in competing and cooperative groups

To gain insights into ecological mechanisms underlying the divide of co-occurring communities between competitive and cooperative types, we compared the metabolic features of the respective member species. We observed that the species present in cooperative communities have fewer metabolic genes (mean 550, sd 65) compared to all the species mapped in the EMP dataset (mean 723, sd 62) (Fig. 2a). We then estimated the minimal nutritional requirements of each species (accordingly to the model simulations, discounting for inorganic compounds). In line with their small metabolic networks, species in cooperative communities have higher nutrient requirements, requiring an average of 10 (sd 1.6) organic compounds, in comparison to the average of 6.8 (sd 1.2), for all EMP species (Fig. 2b). In contrast, species in the competitive communities have, on average, more metabolic genes (mean 919, sd 75) and fewer nutritional requirements (mean 5.0, sd 1.0).

**Figure 2.**
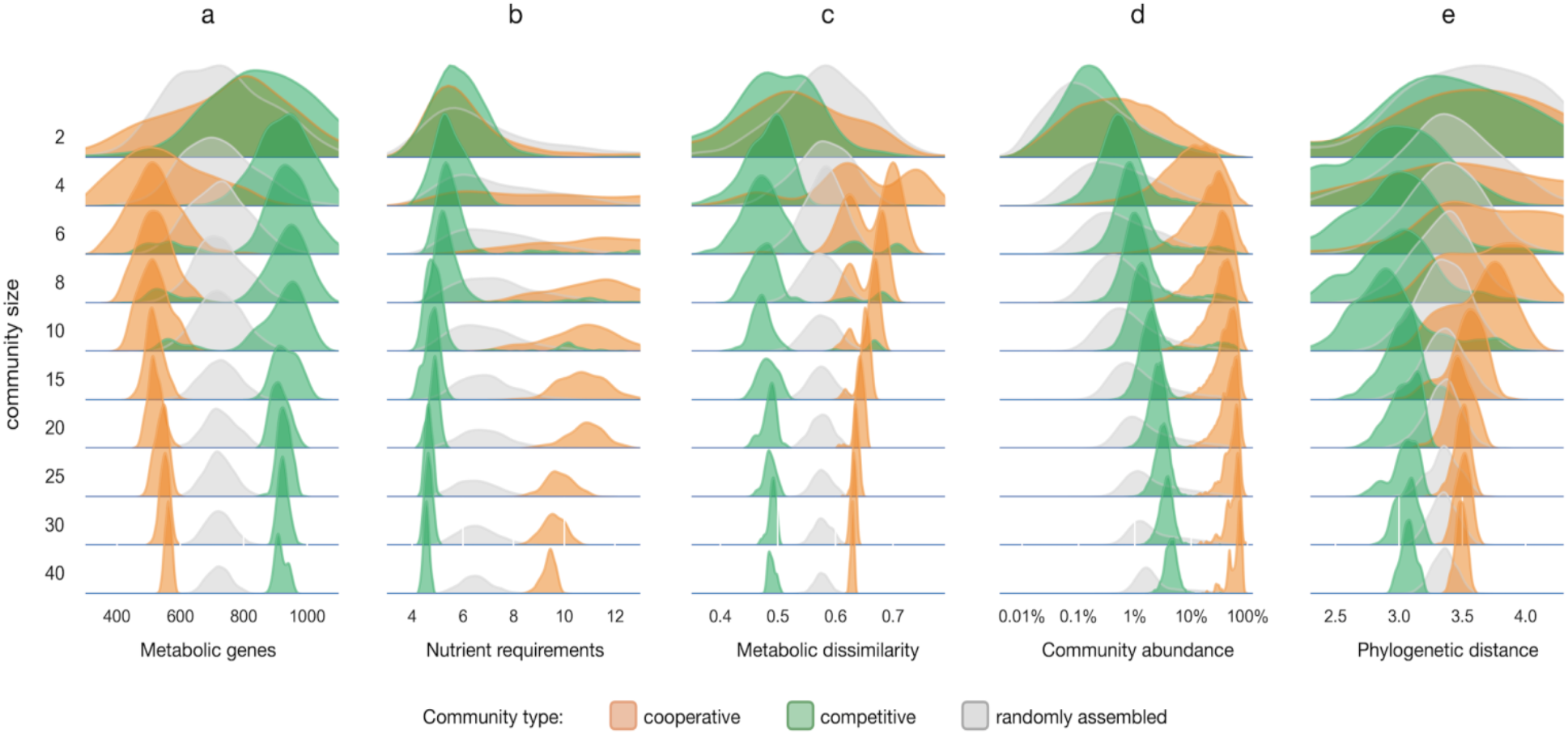
Characteristics of the member species of the cooperative, competitive, and randomly-assembled communities. Shown are the distributions of: a) the number of metabolic genes; b) the number of nutrients (discounting inorganic compounds) required; c) the dissimilarity (Jaccard distance) between the metabolic networks of all species pairs; d) the abundance of community members across all the samples wherein the community occurs (in this case, the random assemblies correspond to random subcommunities of equal size taken from the same samples); e) phylogenetic distance between all pairs of species.

The fact that the cooperative communities have a smaller resource overlap amongst their member species, despite requiring more nutrients, seems puzzling, but can be explained by possible diversification of their nutritional requirements. To test this, we calculated the network dissimilarity within each community (defined as the average Jaccard distance between the metabolic networks of its species). Indeed, the cooperative communities were found to be more dissimilar than expected by chance (Cohen’s d = 1.1, p < 0.001) (Fig. 2c), explaining their lower resource overlap and higher cross-feeding potential. Conversely, the lower dissimilarity in competitive communities (Cohen’s d = −2.0, p < 0.001) was consistent with their high resource overlap.

We next analyzed the nature of the compounds that different communities compete for or exchange within their member species (Supp. Fig. 4). While the cooperative communities mainly require amino acids, the competitive communities showed a more uniform distribution of requirements, including amino acids, carbohydrates, and pyrimidines. Regarding nutrient exchange, cooperative communities showed a three-fold higher propensity for amino acid crossfeeding than the competitive communities (Welch’s t-test: p < 0.001).

To what extent do the cooperative or competitive metabolic strategies of a community influence the fitness of its members? To answer this, we calculated the total (relative) abundance of the species that participate in co-occurring communities and compared it with the abundance of random subsets with the same number of species (Fig. 2d). We find that the species forming cooperative communities are highly abundant, representing a large fraction of the total biomass in each sample (in the range of 10-100%, median 21.6%). In stark contrast, species participating in competitive communities are only slightly more abundant (median 1.2%) than expected by chance (median 0.5%). Since a given species can be part of multiple co-occurring communities, we analyzed how the fitness of an individual species (in terms of its relative abundance) is related to the total number of co-occurring partners present in each sample (Supp. Fig. 5). While for the competitive species the number of partners present does not seem to influence their abundance (Spearman’s r = −0.02, p < 0.05), the species participating in cooperative communities have a higher abundance when more cooperative members are also present in the same samples (Spearman’s r = 0.28, p < 0.001). Membership of a cross-feeding community thus seems to carry a substantial benefit in terms of increased fitness.

### Habitat preferences of cooperative and competitive communities

The evolution of metabolic competition and cooperation is expected to be driven by nutrient availability in the habitat. We tested this by using the EMP ontology describing 17 types of habitats (9 free-living and 8 host-associated)^16^. When counting the number of distinct habitats in which all members of a given community co-occur, we observed a striking difference between competitive and cooperative communities (Fig. 3). The competitive communities are mostly present in free-living environments, with over two-thirds of the respective samples coming from studies in soil diversity^27–29^. On the other hand, the cooperative communities are present both in free-living and host-associated habitats, with the respective samples coming from varied sources, including studies of the human microbiome and studies of microbes present in indoor environments and wastewater treatment^30–33^. Interestingly, we noted several indoor environment samples where competitive and cooperative communities co-exist. These samples come from the Home Microbiome Project (HMP), a study that tracked the microbiome of 18 individuals and 4 pets at multiple body sites, as well as multiple indoor surfaces during 6 weeks. These samples thus likely represent encounters of bacteria from soil, pets, and humans during daily life activity.

**Figure 3.**
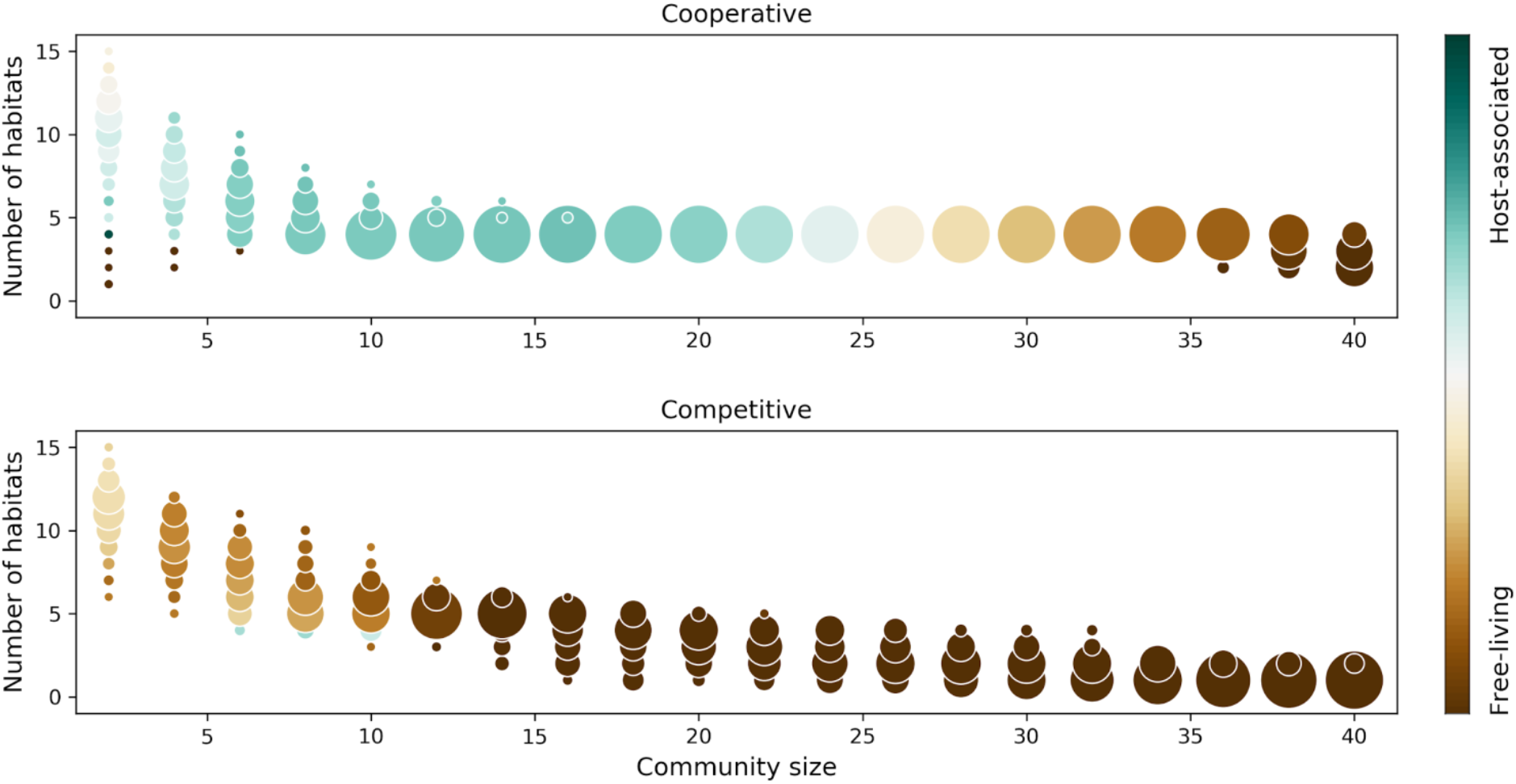
Habitat preferences of co-occurring communities. Circle position indicates the total number of habitats (according to EMPO level 3) for given community size, and the circle size indicates the fraction of communities that live in that number of habitats (out of 1000 computed communities per size). The circle color indicates the ratio between host-associated and free-living habitats.

The high habitat diversity of the cooperative communities supports their advantage as a group, enabling movement between different environments as largely self-sufficient modules. However, to maintain their function, and exchange all required metabolic precursors in suitable amounts, these modules would need to maintain a stable composition in terms of the relative abundance of its members. Therefore, we queried the temporal stability of cooperative and competitive communities in the HMP samples, using two different metrics (see Methods): individual stability (i.e. how stable is the abundance of each species over time), and group stability (i.e. how stable is the relative abundance between community members over time). Supporting our hypothesis, the cooperative communities appear to be more stable than expected by chance (Supp. Fig. 6), both in terms of individual stability (lower coefficient of variation in the abundance of each species, Cohen’s d = −3.5, p < 0.001) and group stability (higher similarity of abundance profiles, Cohen’s d = 3.2, p < 0.001). Competitive communities, on the other hand, showed no coherent trend. Together, the habitat preference analysis brings forward cooperative communities as functionally coupled modules that can successfully migrate between different environments.

### Phylogeny and evolution

We next investigated the role played by gene loss (or gain) by the member species. To address this, we reconstructed a phylogenetic tree (using 40 universal marker genes) for all species in the EMP dataset (see Methods) and calculated the average phylogenetic distance within the members of cooperative and competitive communities. Members of both competitive and cooperative groups are observed across the four main phyla (Fig. 4a), indicating that this polarization is a broadly distributed phenomenon. Also, we observe that the species participating in cooperative communities are phylogenetically more distant than expected by chance (Cohen’s d = 0.3, p < 0.001), whereas competitive communities are closer to each other (Cohen’s d = −0.9, p < 0.001) (Fig. 2e). This agrees with metabolic dissimilarity as one of the distinguishing features between cooperative and competitive communities (Fig. 2c).

**Figure 4.**
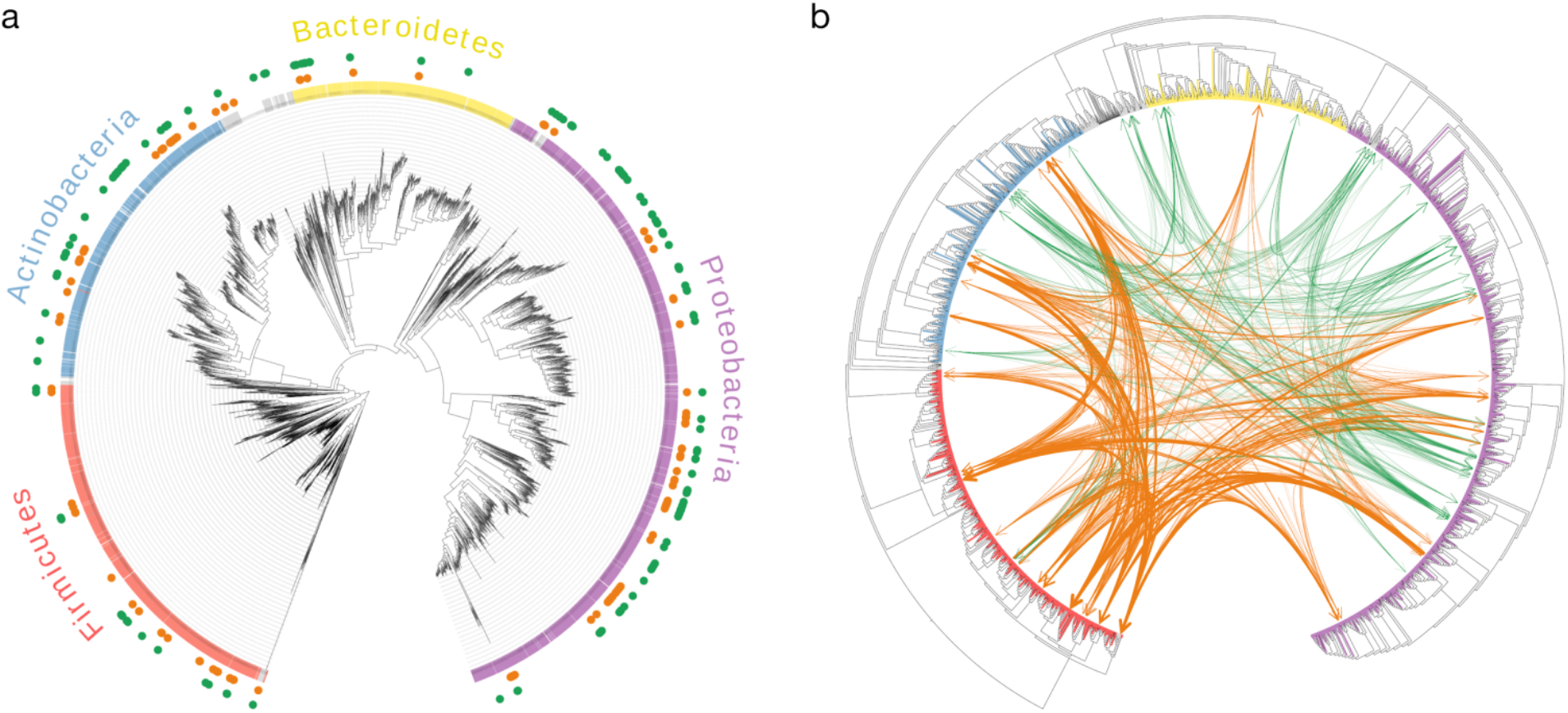
Phylogenetic trees for all the species in the EMP dataset that could be mapped to reference genomes. a) distribution of competitive (green) and cooperative species (orange) across the four main phyla; b) predicted cross-feeding interactions between the 50 most frequently occurring species in competitive (green) and cooperative (orange) communities. The edge width represents the SMETANA score for each interaction (indicating the frequency and total number of exchanged compounds).

To gain insights into the connection between phylogenetic relatedness and inter-species metabolic dependencies, we calculated cross-feeding scores between the 50 most frequently cooccurring species within each community type (see Methods). As expected, we observe stronger interactions between species in cooperative communities (Fig. 4b). Notably, inter-phylum interactions seem to be the rule rather than the exception (2.8-fold more frequent than intraphylum interactions). Cross-feeding interactions are predominant in Firmicutes with both Actinobacteria and Proteobacteria. In agreement, interactions between these phyla have been experimentally observed^34^ and reported in systematic reviews of microbial interactions^35,36^.

Our data shows that cross-feeding of amino acids is the most prevalent type of interaction in cooperative communities. This agrees with previous experimental observations reporting amino acid exchange as one of the main drivers of community interactions^12,35,37^. Moreover, engineered complementary amino acid auxotrophies have been shown to enable the stable assembly of synthetic microbial communities^38,39^. Spontaneous acquisition of amino acid auxotrophies and cross-feeding has also been observed during laboratory evolution of *E. coli*^40^. However, there is still limited evidence for the co-evolution of amino acid exchanges within multi-species consortia *in natura*^41^. Do complementary auxotrophies precede community assembly or are they a consequence of species co-evolution?

To address this question, we assessed whether the amino acid auxotrophies in the cooperative species (i.e. the members of the cooperative communities) have been acquired after speciation or rather inherited from an ancestral species. For a reliable assessment, we used two complementary approaches. One is based on taxonomy (fraction of auxotrophic species within the same genus), and the other based on phylogeny (ancestral state reconstruction along the phylogenetic tree). While a majority of amino acid auxotrophies (~90%) seem to have been inherited (Supp. Fig. 7), we also observe a few cases (12 in total) indicative of recent auxotrophy acquisition. The latter are most frequent for proline (*G. haemolysans, L. hominis, L. inners)* and methionine (*A. tetradius, M. luteus, R. dentocariosa*). Thus, both the assembly of species with pre-existing auxotrophies and the subsequent gene loss appear to have contributed to the establishment of natural communities.

### Response to perturbations

We next asked whether the distinct metabolic characteristics of the cooperative and competitive communities manifest in differential response to abiotic and biotic perturbations. To answer this, we created 100 community models of each type (with 10 members per community, randomly sampled from the most frequently co-occurring species, see Methods) and simulated their response to abiotic (changes in nutrient availability) and biotic perturbations (introduction of a foreign, i.e. non-member, species) (Fig. 5a–b). In particular, we analyzed how the species interaction network is affected by these perturbations. We define sensitivity as a measure of network reconfiguration upon perturbation (see Methods), with lower sensitivity values being an indicator of community resilience.

**Figure 5.**
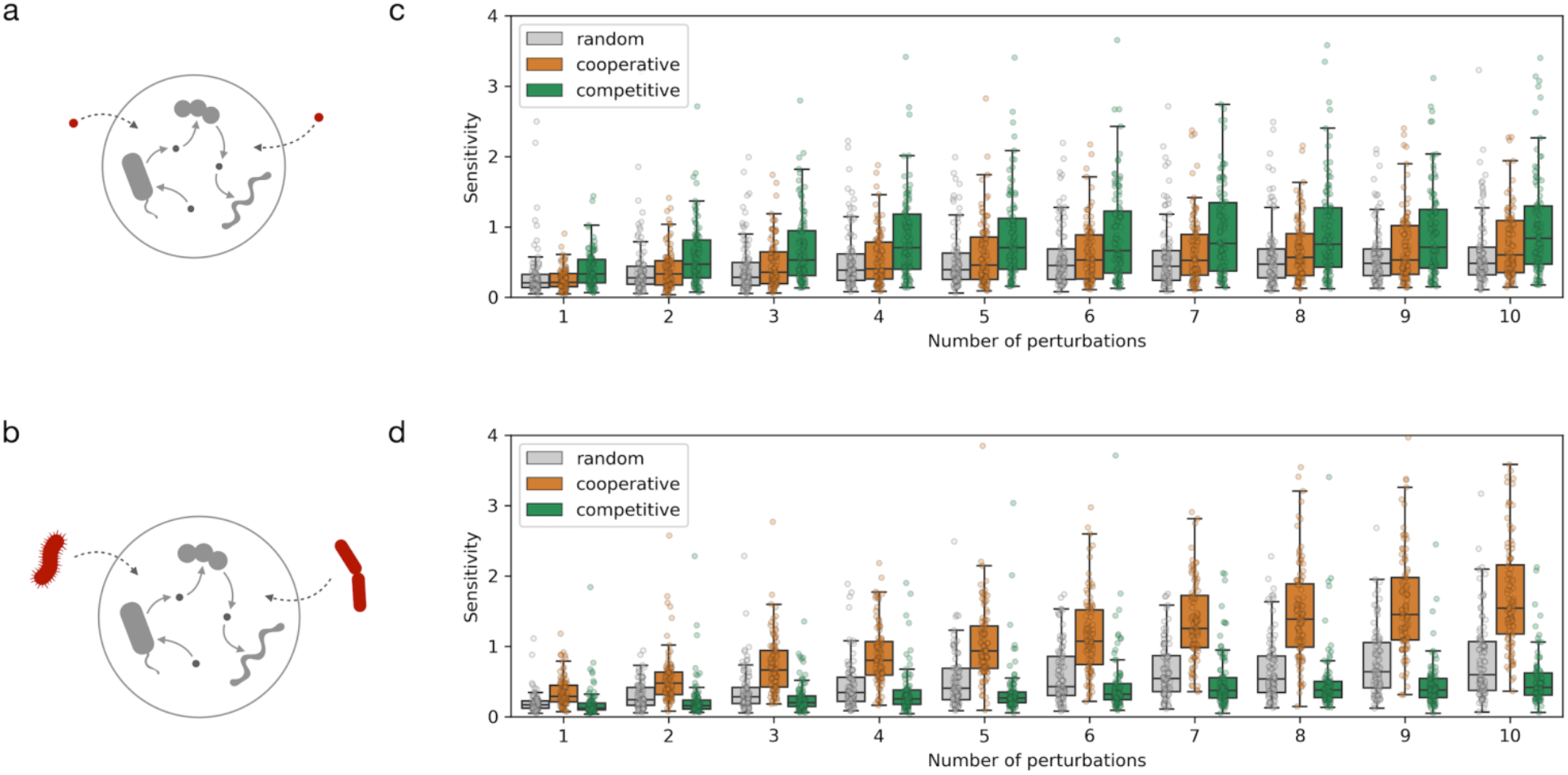
Response of communities to perturbations. a) schematic illustration of abiotic perturbations, changes in medium composition, simulated herein; b) schematic illustration of biotic perturbations, introduction of foreign species in the community, simulated herein; c-d) Simulation results, in terms of sensitivity measure, of cooperative, competitive, and control (randomly-assembled) communities as a function of the total number of simultaneous abiotic (c) and biotic (d) perturbations.

The competitive communities were found to be more sensitive to abiotic perturbations than either cooperative or randomly-assembled communities (Fig. 5c). This was indeed expected as the former are mostly driven by competition for shared resources, and the introduction of different nutrients can result in niche expansion. On the other hand, competitive communities seem very robust to biotic perturbations and less likely to be invaded by foreign species (Fig. 5d). In agreement with these results, Goldford et al have recently shown that the assembly of plant and soil-derived communities (the kind of habitat where we find competitive communities to be prevalent) is primarily determined by the carbon source, and has little influence from the initial species diversity in the inoculum^42^.

Cooperative communities, on the other hand, seem robust to abiotic perturbations, but quite sensitive to the introduction of foreign species (Fig. 5d). The magnitude of the response appears to increase with the number of species introduced. This can be explained by the fact that these communities have multiple cross-feeding interactions, which can be “intercepted” by the invading species, leading to the rewiring of the species network. To test this hypothesis, we analyzed the species invasion pattern in a recent study by Zmora et al, wherein a host-specific response to colonization by probiotics was observed, identifying colonization permissive and colonization resistant individuals^43^. In light of our results, we hypothesized that the microbiomes of permissive and resistant individuals will be characterized, respectively, by the presence of cooperative and competitive species. Confirming this, the individuals permissive to colonization display an increased presence of cooperative species along the lower gastrointestinal (LGI) tract, both in terms of total number of species per location (Wilcoxon signed-rank test, p = 0.017) and their relative abundance (Wilcoxon signed-rank test, p = 0.018) (Supp. Fig. 8). Notably, this difference is more striking in the early LGI track (terminal ileum, cecum, ascending colon). Supporting our hypothesis, the presence of cooperative species appears to facilitate colonization by probiotic species.

## Discussion

The polarization of co-occurring microbial communities into competitive and cooperative groups and its spread across the phylogenetic tree indicates two different, habitat-driven, evolutionary paths in community assembly: the one followed by competitive communities in accordance with the red queen hypothesis^37^, and the other followed by cooperative communities conforming to the black queen hypothesis^38^. Both the red queen and the black queen theories thus seem to be operating in the community assembly *in natura*, reflecting two extremes in the trade-off between competition and cooperation. Our analysis brings forward the abiotic habitat and the evolutionary gene loss as the main drivers determining whether a competitive or cooperative community will be established.

The competitive red queen species are generally restricted to free-living habitats wherein the resources are likely to be more scarce making competition more prevalent. In contrast, the nutritional richness of the host-associated habitats seems to support the more cooperative black queen species, which exhibit complementary auxotrophies, in part resulting from gene loss. This adaptation not only confers a fitness advantage but is also likely to facilitate the survival of these species during migration between the hosts and the external environment as a highly self-sufficient group. The generally higher abundance and diverse habitat occupation of the cooperative groups point to the advantages offered by the division of metabolic labor in the group, and the consequent independence from the environment.

The existence of two community types with contrasting metabolic make-up and habitat preference means that the strategies to modulate or re-engineer these communities also need to be separately tailored. Our *in silico* results, with support from previously published experimental data, show that the competitive communities are more malleable through abiotic perturbations, whereas the cooperative communities are more malleable through biotic perturbations. These findings could, in future, be further refined to consider metagenome-assembled genomes^44^, improving coverage at species and strain levels, and by accounting for viral^45^ and fungal^46^ interactions. Altogether, we conclude that devising intervention strategies tailored to communities according to their position in the competition-cooperation landscape would be key to the modulation of complex microbial ecosystems.

## Methods

### Mapping OTUs to reference genomes

The EMP dataset provides 16S tags with multiple lengths. The abundance table with the longest reads (150 bp) was downloaded from the EMP portal (http://www.earthmicrobiome.org). All reference/representative bacterial genomes were downloaded from NCBI RefSeq (release 84). The 16S tags from the EMP data were mapped to those extracted from the reference genomes using *diamond* with a 97% identity threshold and a 95% alignment coverage. If multiple genomes were found, the ones with the highest alignment identity and length were selected. As expected, a large fraction of OTUs did not have a matching assembled genome (and some OTUs matched the same genomes). Overall, the diversity in each sample is reduced by almost an order of magnitude (from an average of 990 OTUs per sample to an average of 159 genomes) (Supp. Fig. 1a). Nevertheless, we observe an enrichment regarding species prevalence (7-fold increase, Welch’s t-test: p < 0.001) and abundance (2.5-fold increase, Welch’s t-test: p < 0.001) (Supp. Fig. 1b,c), indicating that the unmapped OTUs are associated with less prevalent and less abundant species. This was also reflected when we compared the fraction of OTUs covered per sample (mean 20.8%) to their relative abundance (mean 40.6%) (Supp. Fig. 1d).

### Computing co-occurrence

We computed higher-order co-occurrence using an iterative algorithm that begins with species pairs and gradually computes co-occurring groups of larger sizes. At the beginning of each iteration, all combinations of species are evaluated for co-occurrence by counting the total number of samples in which they co-occur. We calculate the number of co-occurring observations expected by chance using a binomial distribution and the probability of observing each species individually. We also calculate FDR-corrected p-values (q = 0.05), and select all species combinations that: 1) co-occur in at least 10 samples; 2) co-occur at least twice more than expected by chance; 3) pass the FDR-correction test. The 1000 most frequently co-occurring groups of species are selected as the best solutions for the current group size. In the next iteration, larger sets are created by extending all groups with a new element from the complete set of species in a combinatorial manner. To cope with the combinatorial explosion, at each iteration we only propagate a population of 10,000 solutions to the next iteration. This population is randomly selected using roulette wheel selection with a probability proportional to the co-occurrence frequency. The presence/absence of a species in any sample is evaluated with a relative abundance cutoff. In particular, using cut-off values of 0.1% and 0.01% originated the two different community types analyzed in this study. We tried higher and lower cut-off values, without any observable differences in the results (Supp. Fig. 2).

This method is implemented as a standalone python package, HiOrCo, openly available at https://github.com/cdanielmachado/hiorco.

### Community simulation

All simulations were performed using SMETANA v1.0. The different scores computed with SMETANA used in this study (such as MIP and MRO) are described in its original publication^23^. The tool is implemented as a standalone python package, and is openly available at https://github.com/cdanielmachado/smetana.

### Phylogenetic analysis

A maximum likelihood-approximate phylogenetic tree of 2992 species of Prokaryotes was built using ETE3 toolkit^47^ with JTT model^48^ by aligning protein sequences of the 40 conserved universal marker genes^49,50^ with default parameters in the ClustalOmega aligner^51^ and FastTree2 treebuilder^52^. A cophenetic distance matrix was constructed from the tree using the ape package^53^ in R (v3.4.4). The phylogenetic trees were visualized and exported with iTOL^54^.

To estimate the ancestral state of amino acid auxotrophies, we first calculated auxotrophies for all reference species using genome-scale metabolic models. We then used the *make.simmap* function from the phytools^55^ library in R (v.3.4.4) with 100 stochastic character mappings followed by the *describe.simmap* function to obtain posterior probabilities of auxotrophic ancestral state (considered as a discrete trait). This method relies on stochastic character mapping that is sampled from a Markov chain Monte Carlo Bayesian posterior probabilities distribution^56^.

### Simulating community response to perturbations

For each of the two types of communities (competitive and cooperative), we selected the 50 most representative species (i.e., those that are most frequently present in all co-occurring communities), and used them to randomly generate 100 communities of 10 species each. Each community was subject to multiple random perturbations with N perturbation elements (up to 10). In the abiotic case, the perturbations consisted of 100 random perturbations per community adding N additional nutrients to the growth medium. The biotic case consisted of 10 random perturbations per community, introducing N foreign species. The sensitivity to perturbations is calculated as follows:

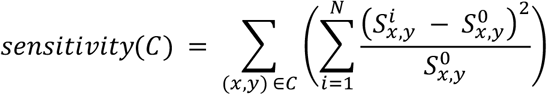

where *N* is the number of perturbations, (*x,y*) is a pair of species in community *C*, and *S_x,y_* is the SMETANA score for cross-feeding interactions between *x* and *y*, defined as:

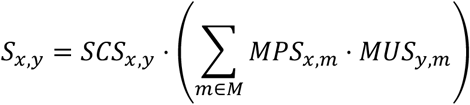

where *SCS* (species coupling score), *MPS* (metabolite production score) and *MUS* (metabolite uptake score) are calculated as defined in Zelezniak et al^23^, and *M* is the complete set of metabolites that can be produced and consumed.

### Community stability analysis

Individual species stability was calculated as the coefficient of variation of the relative species abundance across all time points in a given sample:

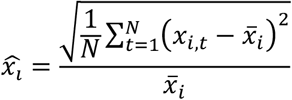

where *x_i,t_* is the relative abundance of species *i* at time point *t* and *N* is the number of time points.

Group stability was calculated as the average cosine distance *D_i,j_* between the time-course profiles of every pair of species in a group and is defined as:

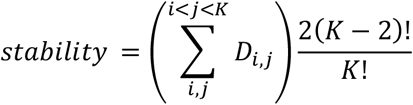

where *K* is the number of species in the group.

For each sample, we computed the stability of the competitive and cooperative subcommunities present in those samples as well as the stability of 100 randomly-assembled subcommunities.

## Acknowledgments

The authors would like to acknowledge Niv Zmora and Eran Elinav for providing data and feedback on the probiotics study, and Sebastian Schmidt for fruitful discussions. This project has received funding from the European Union’s Horizon 2020 research and innovation programme under grant agreement No 686070.

## Author Contributions

D.M. developed the co-occurrence computation method and performed the simulations and data analysis. D.M. and S.A implemented the simulation software. O.M.M. performed the phylogenetic analysis. Y.K. mapped the OTUs to reference genomes. P.B. supervised the phylogenetic analysis. Kaustubh R.P. and Kiran R.P. conceived the study. D.M. and Kiran R.P. wrote the manuscript. All authors read and revised the final manuscript.

## Data and code availability

All the data and code required to generate the results and figures presented in this article are publicly available in the following repository: https://github.com/cdanielmachado/cooccurrence.

## Supplementary Figures

**Supplementary Figure 1:**
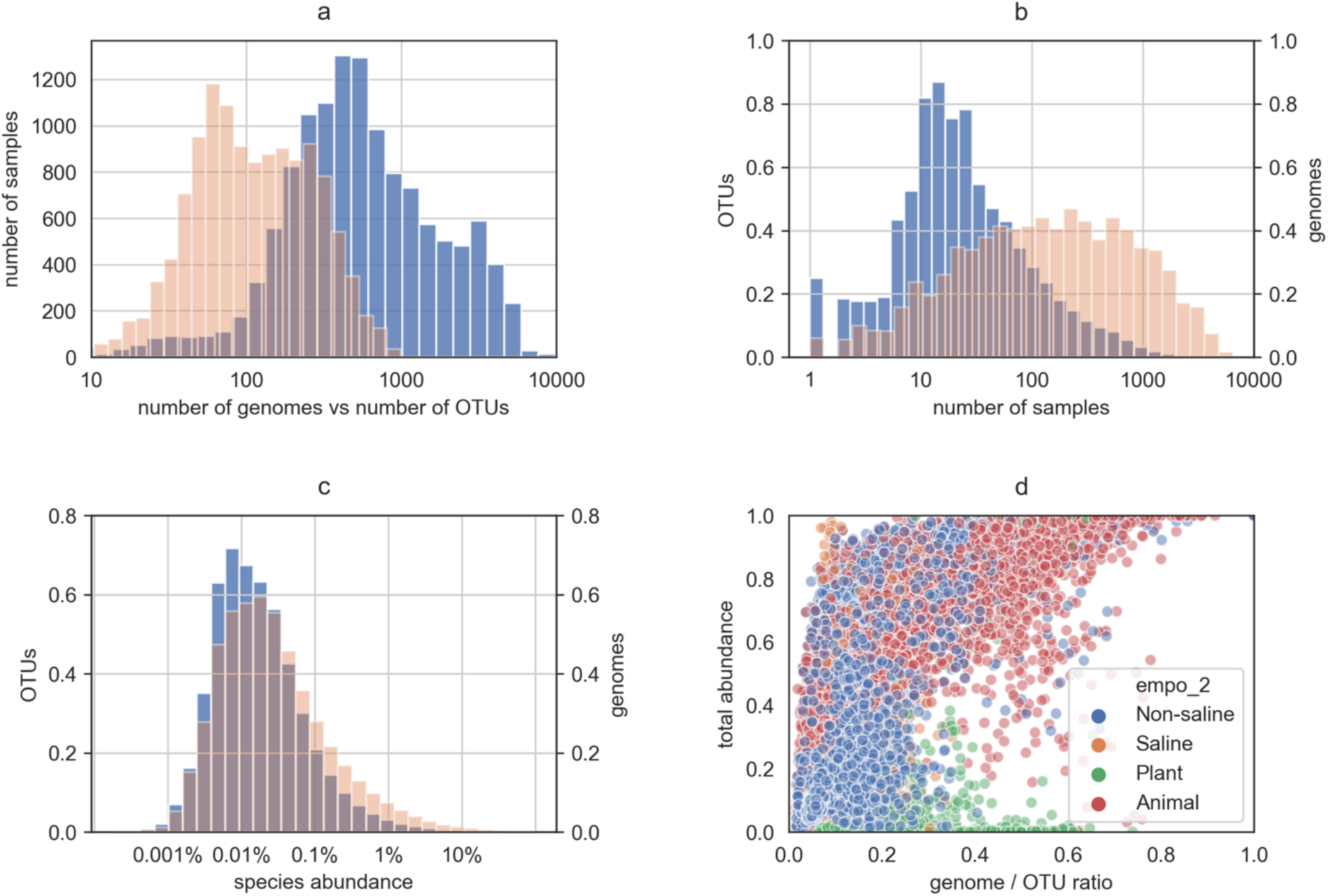
Results for mapping OTUs to reference genomes: a) comparison of sample diversity in terms of OTUs (blue) and genomes (orange); b) comparison of species prevalence in terms of OTUs and genomes across samples; c) species abundance distribution; d) total abundance of each sample that is captured by the mapped genomes in comparison to the ratio of genomes to OTUs.

**Supplementary Figure 2:**
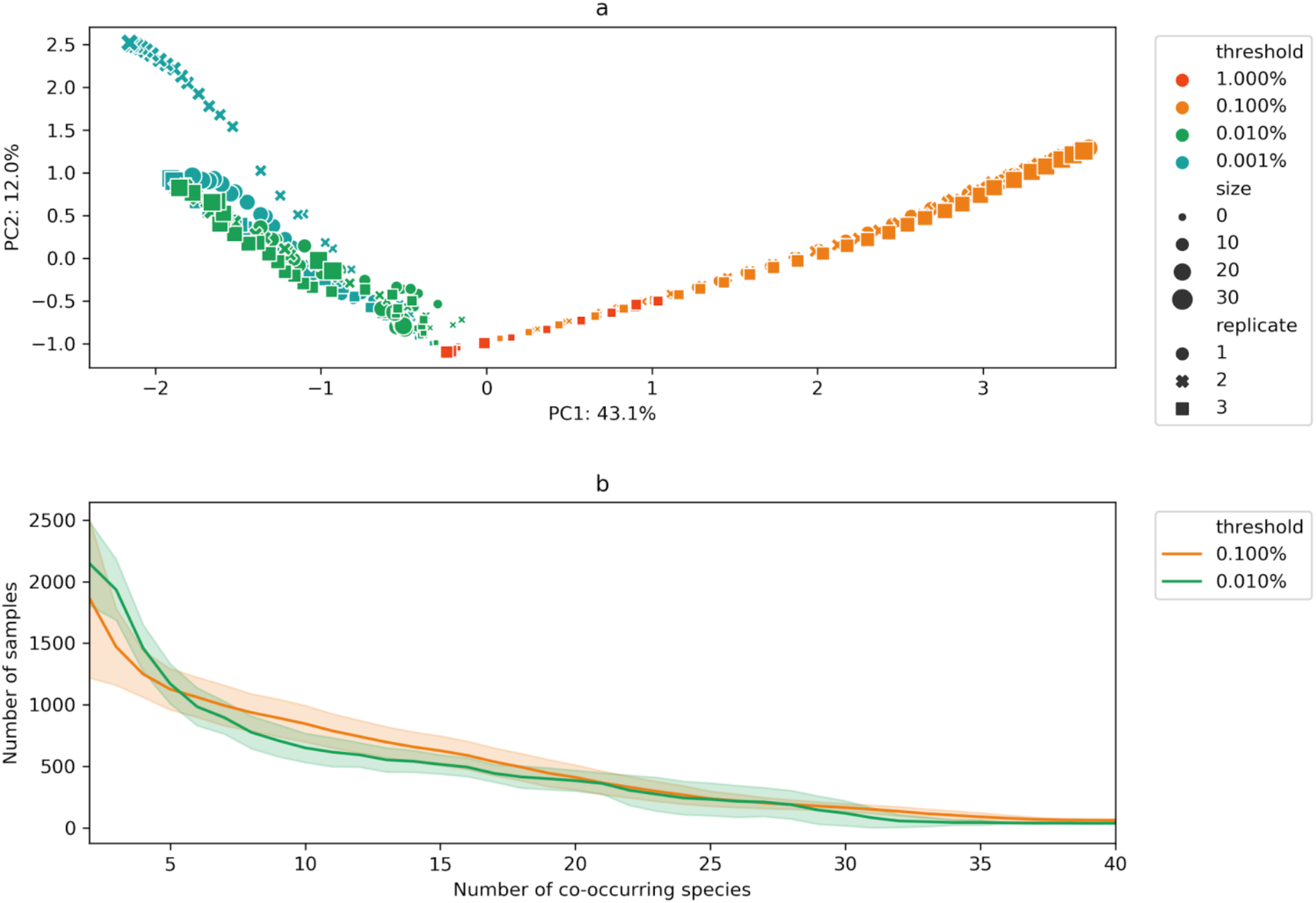
a) Principal component analysis of co-occurring communities computed using different abundance thresholds. Marker size indicates co-occurring community size (up to 30 species) and marker shapes indicates independent runs of the algorithm (3 runs for each threshold); b) Average number of samples where all species in a co-occurring community can be found together as a function of community size (computed for the threshold values used in this work).

**Supplementary Figure 3:**
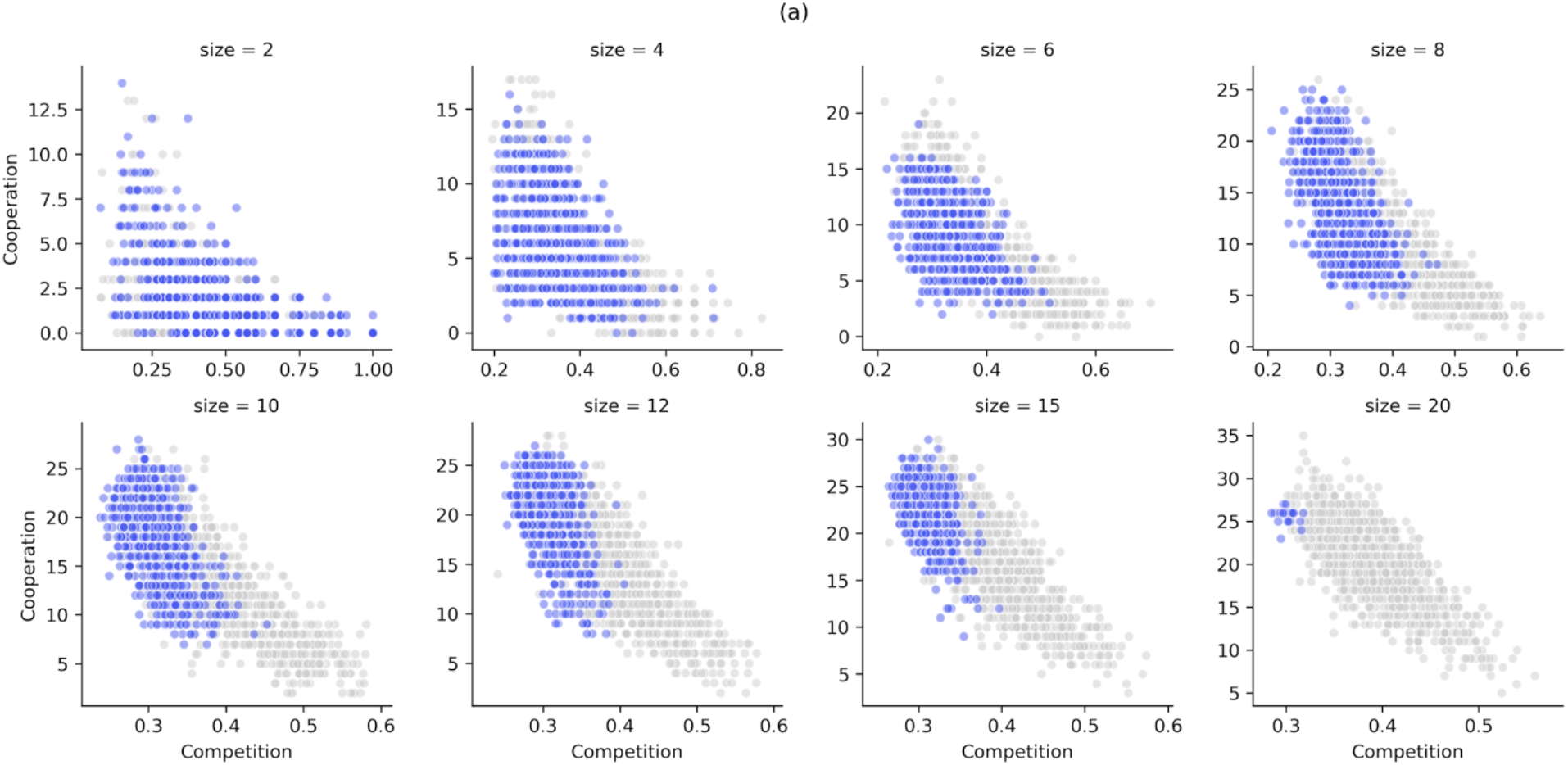
Simulation results for competition (MRO score) and cooperation potential (MIP score) for microbial communities obtained from Chaffron et al. Blue dots represent co-occurring communities of different sizes (up to 1000 per size) and grey dots represent randomly-assembled communities of similar size (1000 communities per size).

**Supplementary Figure 4:**
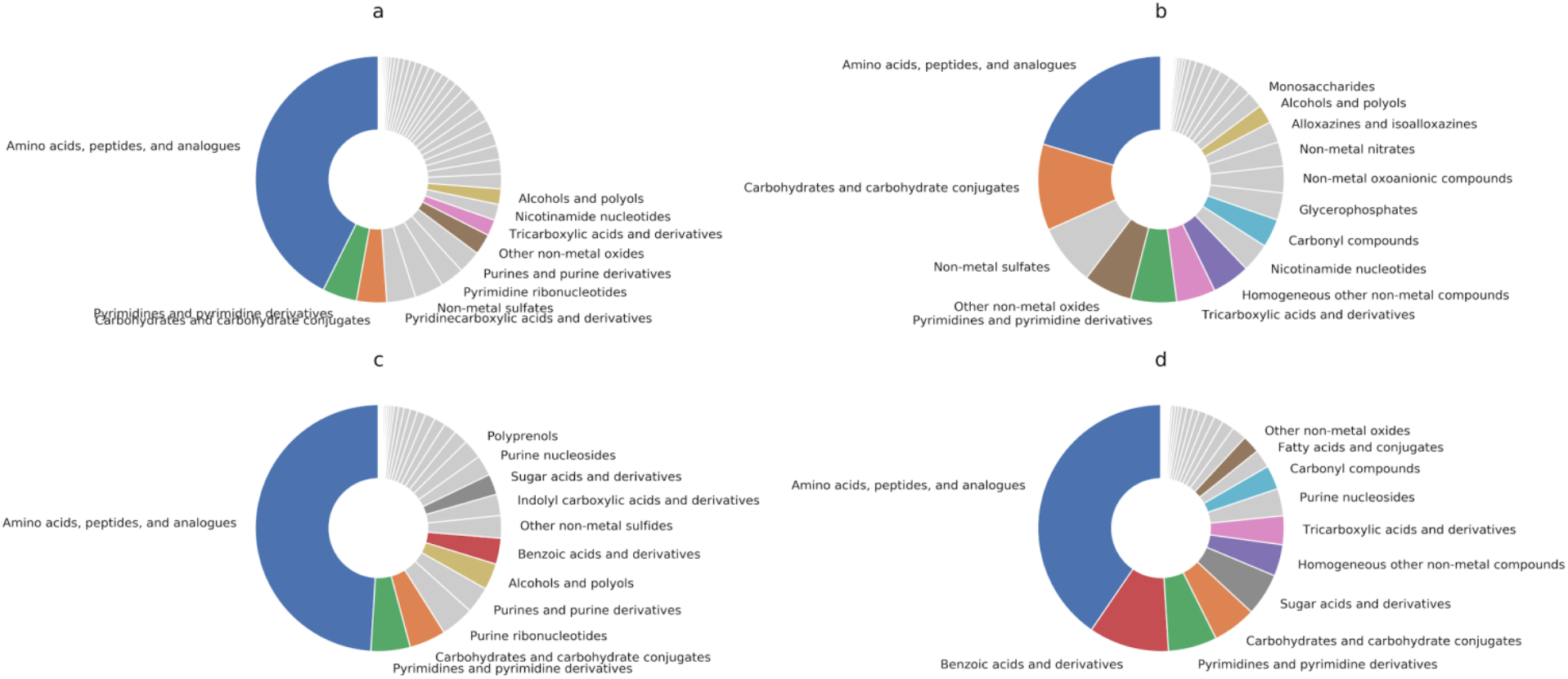
Resource distribution: a) compounds competed for in cooperative communities; b) compounds competed for in competitive communities; c) cross-fed compounds in cooperative communities; d) cross-fed compounds in competitive communities. Compound classification according to the Human Metabolome Database (HMDB).

**Supplementary Figure 5:**
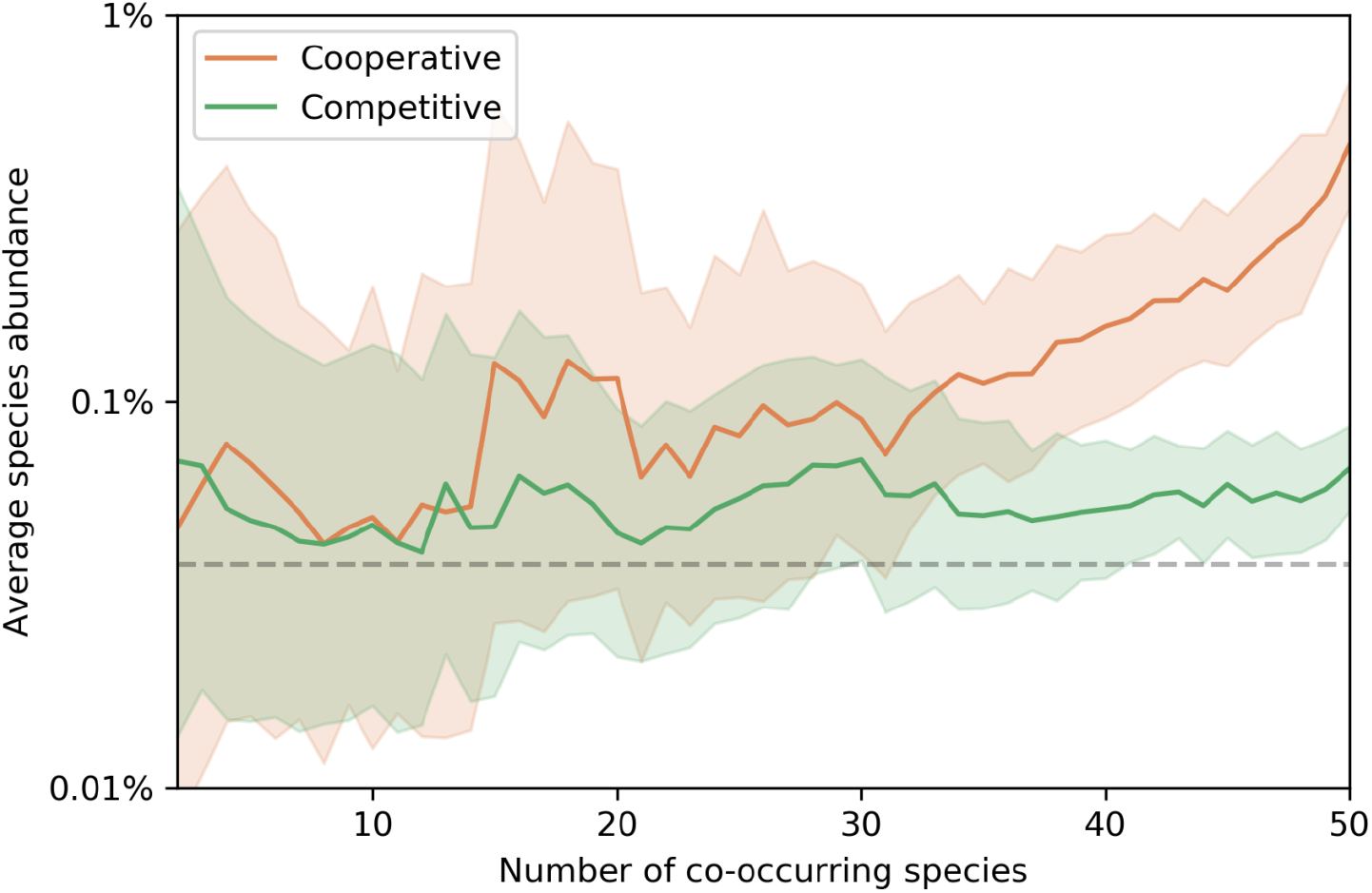
Average species abundance of co-occurring species as a function of the total number of co-occurring species present in each sample. The colored line denotes the average abundance for each type of community, the shadowed area indicates standard deviation, and the dashed grey line indicates the average species abundance across all species and samples. In all cases the average is calculated as the mean relative abundance value in log-space (i.e. the geometric mean of the relative abundances).

**Supplementary Figure 6:**
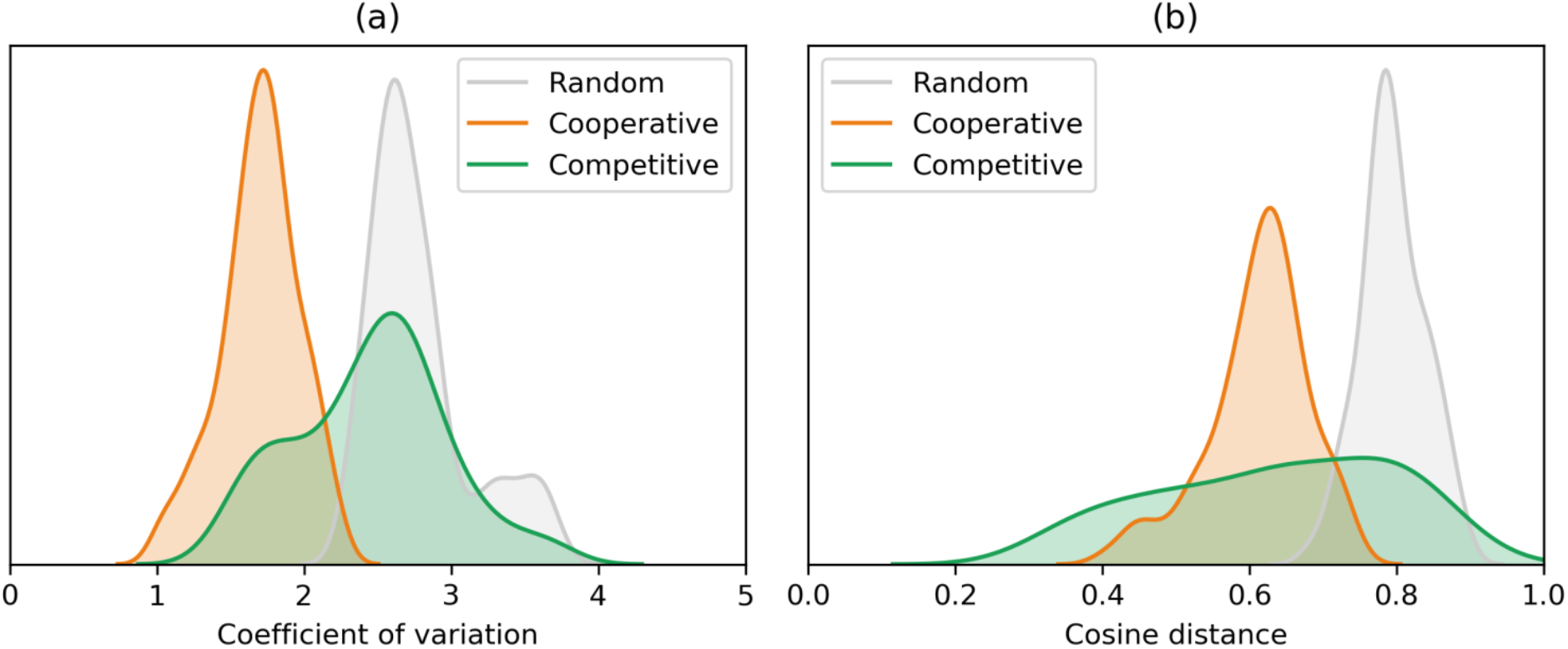
Community stability measured in terms of: a) individual stability (lower coefficient of variation per species indicates higher stability); b) group stability (lower cosine distance indicates higher covariation of species abundance within each community).

**Supplementary Figure 7:**
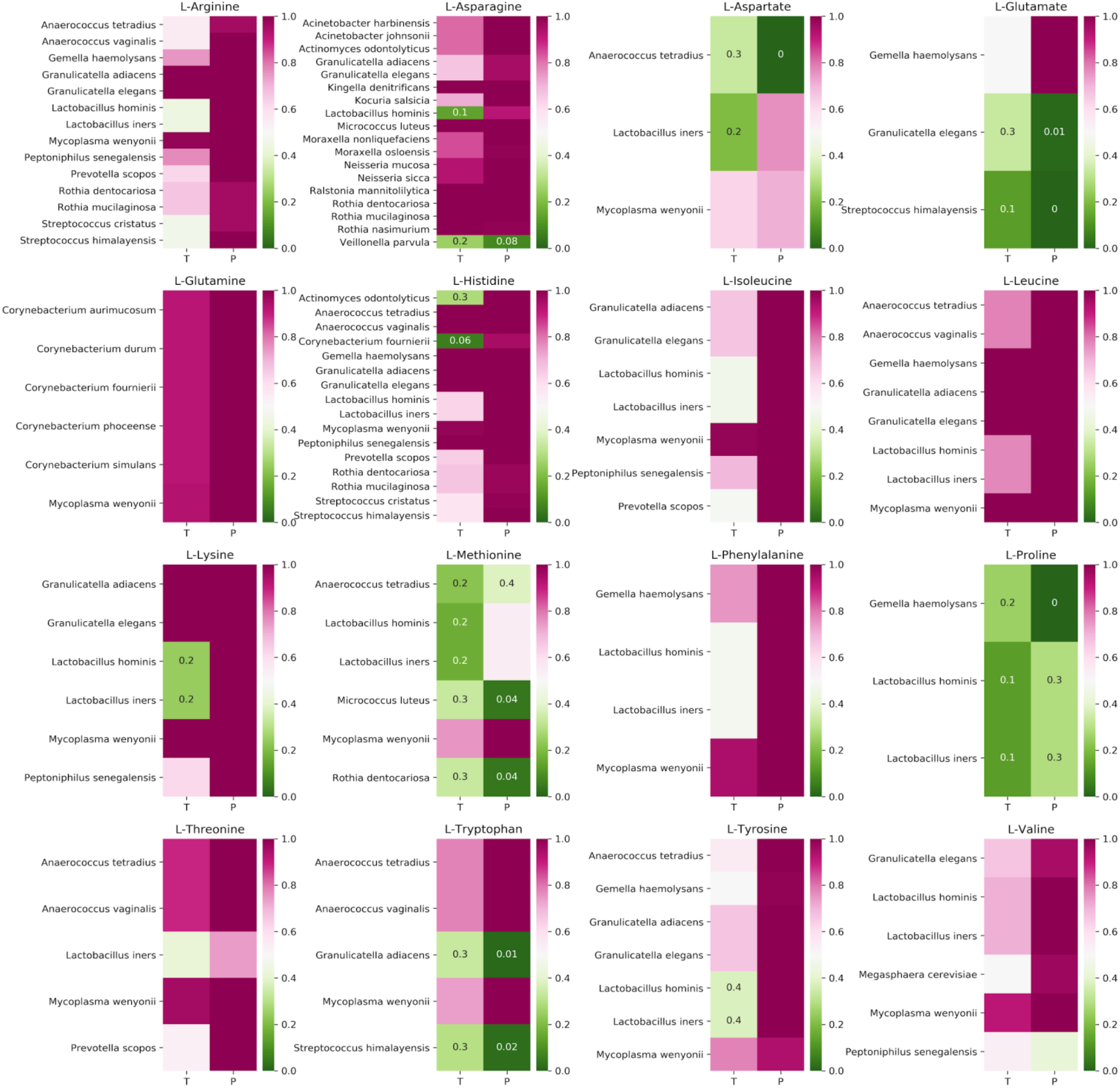
Analysing the recent acquisition of amino acid auxotrophies using two complementary approaches: taxonomy based (T), measuring the fraction of auxotrophic species at genus level; phylogeny based (P), estimating the probability of the auxotrophy being present in the most recent ancestor of the species. Green color in both columns is indicative of a consensus.

**Supplementary Figure 8:**
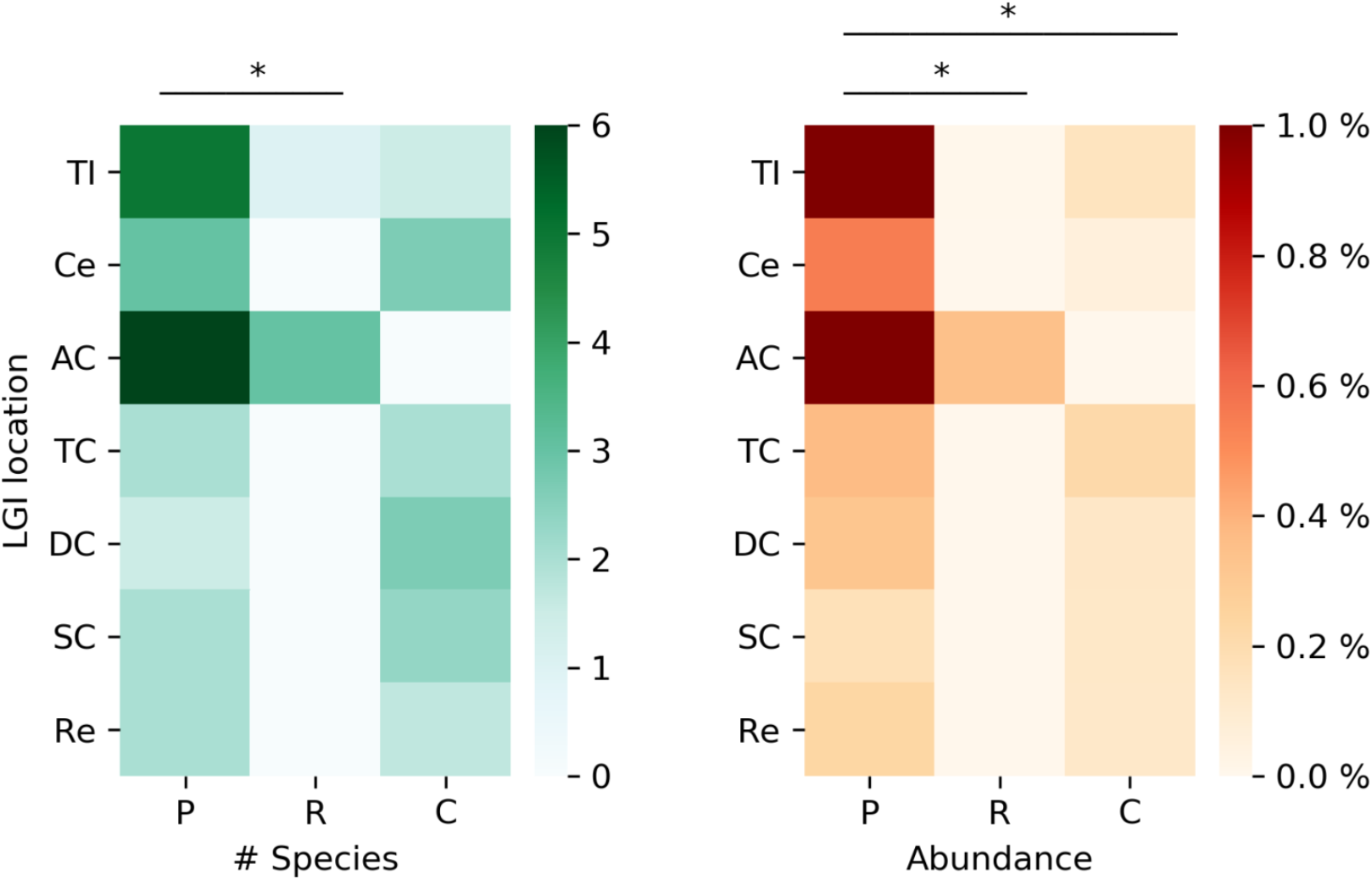
Presence of cooperative species in the lower gastrointestinal (LGI) tract of patients permissive to probiotic colonization (P), patients resistant to colonization (R) and control patients (C). Asterisks indicate significance of Wilcoxon signed-rank test (p < 0.05).

